# High Genetic Potential for Proteolytic Decomposition in Northern Peatland Ecosystems

**DOI:** 10.1101/375428

**Authors:** Emily B. Graham, Fan Yang, Sheryl Bell, Kirsten S. Hofmockel

## Abstract

**Abstract:** Nitrogen (N) is a scarce nutrient commonly limiting primary productivity. Microbial decomposition of complex carbon (C) into small organic molecules (e.g., free amino acids) has been suggested to supplement biologically-fixed N in high latitude peatlands. We evaluated the microbial (fungal, bacterial, and archaeal) genetic potential for organic N depolymerization in peatlands at Marcell Experimental Forest (MEF) in northern Minnesota. We used guided gene assembly to examine the abundance and diversity of protease genes; and further compared to those of N-fixing (*nifH*) genes in shotgun metagenomic data collected across depth at two distinct peatland environments (bogs and fens). Microbial proteases greatly outnumbered *nifH* genes with the most abundant gene families (archaeal M1 and bacterial Trypsin) each containing more sequences than all sequences attributed to *nifH*. Bacterial protease gene assemblies were diverse and abundant across depth profiles, indicating a role for bacteria in releasing free amino acids from peptides through depolymerization of older organic material and contrasting the paradigm of fungal dominance in depolymerization in forest soils. Although protease gene assemblies for fungi were much less abundant overall than for bacteria, fungi were prevalent in surface samples and therefore may be vital in degrading large soil polymers from fresh plant inputs during early stage of depolymerization. In total, we demonstrate that depolymerization enzymes from a diverse suite of microorganisms, including understudied bacterial and archaeal lineages, are likely to play a substantial role in C and N cycling within northern peatlands.

**Importance:** Nitrogen (N) is a common limitation on primary productivity, and its source remains unresolved in northern peatlands that are vulnerable to environmental change. Decompositionof complex organic matter into free amino acids has been proposed as an important N source, but the genetic potential of microorganisms mediating this process has not been examined. Such information can elucidate possible responses of northern peatlands to environmental change. We show high genetic potential for microbial production of free amino acids across a range of microbial guilds. In particular, the abundance and diversity of bacterial genes encoding proteolytic activity suggests a predominant role for bacteria in regulating productivity and contrasts a paradigm of fungal dominance of organic N decomposition. Our results expand our current understanding of coupled carbon and nitrogen cycles in north peatlands and indicate that understudied bacterial and archaeal lineages may be central in this ecosystem’s response to environmental change.

## Introduction

Understanding the processes that govern coupled carbon (C) and nutrient dynamics in northern peatlands is critical to predicting future biogeochemical cycles. These ecosystems account for 15-30% of global soil carbon storage (1-3), primarily occurring within layers of partially decomposed plant materials where nitrogen (N) content is low (4, 5). Nitrogen is a critical nutrient regulating primary productivity in many terrestrial ecosystems (6) and can dictate belowground carbon storage through impacts on soil organic matter decomposition (7, 8). Ombrotrophic peatlands are characterized by *Sphagnum* moss that has a comparatively large N requirement (approximately 40-50 kg ha-1 year-1 of N (9-12). Nitrogen fixation historically has been considered to be the primary N source in peatlands (13-18). Yet, previous work has shown that N fixation alone cannot meet peatland N requirements (5, 19) and many studies have demonstrated the importance of organic molecules in fulfilling N demand (20-25). Symbiotic fungi are traditionally associated with organic N acquisition (22, 25), but there is an increasing appreciation for the role of bacteria in this process. Despite these advances, our understanding of the genetic mechanisms mediating N available remains nascent. We address this knowledge gap by exploring the genetic potential of peatland microbiomes to decompose polymeric organic N and subsequently influence peatland C and N cycles.

Depolymerization of proteinaceous organic material is an important pathway for generating bioavailable N in wide range of systems including boreal forests and is often considered a fungal trait (5, 7, 20, 22, 26-28). Depolymerization decomposes polymeric organic material into monomers and amino acids that can be used as C and N sources by soil microorganisms and plants (19, 20, 22, 29, 30). Several studies from terrestrial ecosystems under strong inorganic N-limitation have shown that organic N, and free amino acids in particular, can be used directly by plants (19, 20, 22, 29, 30). Additionally, microorganisms (defined here as bacteria, archaea, and fungi) secrete extracellular proteases into soils to carry out organic matter depolymerization. Proteases are highly diverse and ubiquitous in soil and provide a large proportion of bioavailable N (31, 32). These enzymes catalyze the initial hydrolysis of proteins into smaller organic molecules such as oligopeptides and amino acids that can be subsequently acquired by plants (31)

In peatlands, fungi are considered more important than bacteria or archaea in proteolytic activity and decomposition more generally, particularly within the oxic surface layer (33-35). Symbiotic ectomycorrhizal and ericoid fungi (EEM), which are supplied with C by a host plant, are especially relevant to organic N depolymerization in peatlands through N mining (36-38). EEM have been suggested to acquire N from soil organic matter (19, 39-41) and enable plants to directly compete with free-living microorganisms for N (42-44), and Orwin et al. (45) posited a critical role for EEM in generating microbial N limitation of decomposition by enhancing plant N uptake. This fungal-mediated plant organic N uptake may be particularly important in N-poor boreal ecosystems (19, 20). In these systems, free-living microorganisms should retain amino acids for growth instead of mineralizing organic N (22, 46). However, empirical evidence supporting the notion that fungi dominate proteolytic activity is sparse and primarily derived from correlative studies.

Little is known about the roles various other microorganisms may play in peatland organic N depolymerization, and the genes that encode microbial proteases may provide valuable insight into the coupling of C and N cycles in these systems. Recent work has suggested substantial involvement of bacteria in depolymerization. For example, Lin et al. (34, 47) indicated that bacteria may outcompete fungal communities for plant-derived substrates, including large polymeric molecules. Consistent with this work, Bragina et al. (48) demonstrated that peatland *Sphagnum* moss microbiomes contain a high abundance of genes involved in N cycling and recalcitrant organic matter decomposition. The involvement of archaeal proteases in peatland organic N decomposition remains largely unexplored.

Here, we evaluate microbial proteolytic potential relative to N-fixation within the Marcell Experimental Forest (MEF) by examining the genes encoding a suite of microbial proteases vs. the *nifH* gene that is well-ascribed to N-fixation. Previous work has demonstrated that organic matter cycling differs between hydrologically-defined environments within peatlands (e.g., bogs vs. fens) (34, 49-51). Fungal biomass also typically declines with depth as oxygen and root exudates become depleted in soils. Based on these observations, we tested the following hypotheses: i) surface peatland protease genes are attributed more to fungi than bacteria, with a shift towards bacterial sequences as depth increased; ii) bog environments contain more proteolytic potential than fens due to stronger N-limitation; and iii) the total number and the diversity of protease genes decrease with depth corresponding with decreased organic matter inputs.

## Results

### Assembly overview

Out of 24 gene groups constructed and investigated, we assembled 13 gene groups successfully (Table S1). The assembled groups include three housekeeping genes (*rplB, rpb2*_4, and *rpb2*_7), nine protease gene families (eight of which are extracellular), and *nifH* gene. Approximately 34% of the fully covered contigs were annotated as housekeeping genes, 5% were annotated as *nifH* genes, and 61% were annotated as protease genes. Among all metagenomic reads mapped to the annotated fully covered contigs, approximately 83.5% were bacterial, 16% were archaeal, and 0.5% were fungal (Table 1).

**Table 1.**
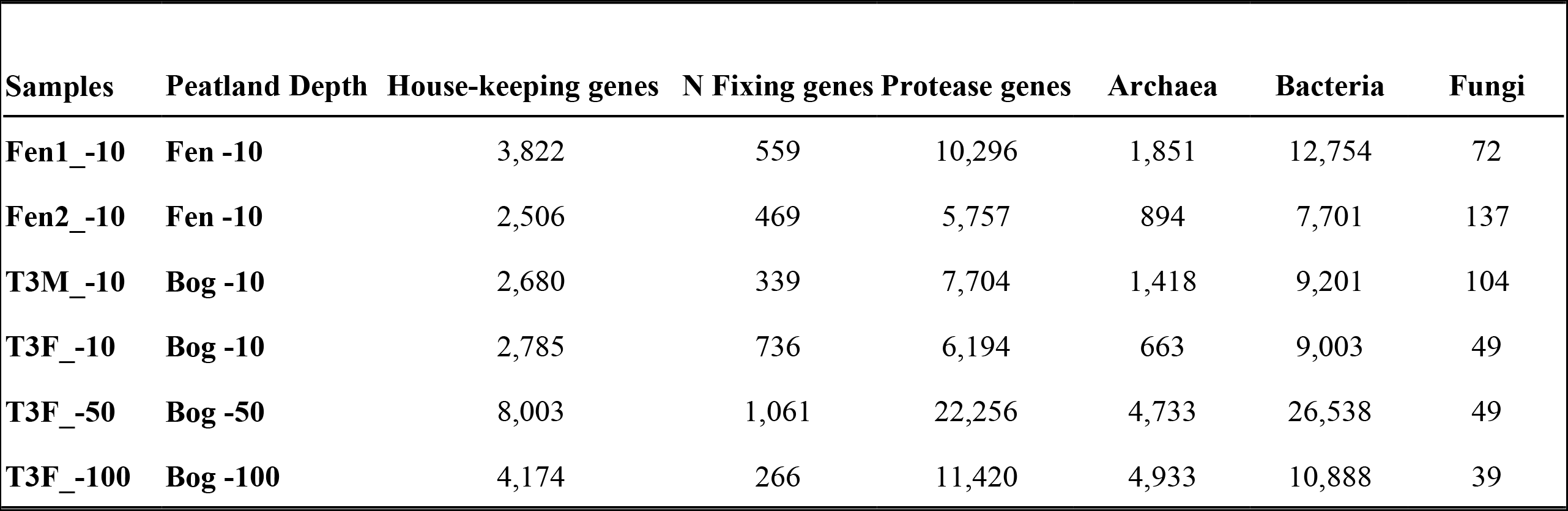
The distribution of mapped reads mapped to contigs identified as housekeeping genes, N fixation genes, and protease genes. The distribution of housekeeping genes among archaea, bacteria, and fungi is also provided.

| Samples | Peatland Depth | House-keeping genes | N Fixing genes | Protease genes | Archaea | Bacteria | Fungi |
| --- | --- | --- | --- | --- | --- | --- | --- |
| **Fen1_-10** | **Fen -10** | 3,822 | 559 | 10,296 | 1,851 | 12,754 | 72 |
| **Fen2_-10** | **Fen -10** | 2,506 | 469 | 5,757 | 894 | 7,701 | 137 |
| **T3M_-10** | **Bog -10** | 2,680 | 339 | 7,704 | 1,418 | 9,201 | 104 |
| **T3F_-10** | **Bog -10** | 2,785 | 736 | 6,194 | 663 | 9,003 | 49 |
| **T3F_-50** | **Bog -50** | 8,003 | 1,061 | 22,256 | 4,733 | 26,538 | 49 |
| **T3F_-100** | **Bog -100** | 4,174 | 266 | 11,420 | 4,933 | 10,888 | 39 |

### Overall gene stratification

The standardized abundance of bacterial genes was similar across sample depth profile, whereas the standardized fungal gene abundance decreased and archaeal genes increased along the sampling depth (Figure 1). Archaeal N-acquisition genes in acrotelm samples were approximately 100 fold more abundant than archaeal housekeeping genes. Fungal protease encoding genes were consistently detected through the depth profile but fungal housekeeping genes were only detected in acrotelm (0-10 cm) samples.

**Figure 1.**
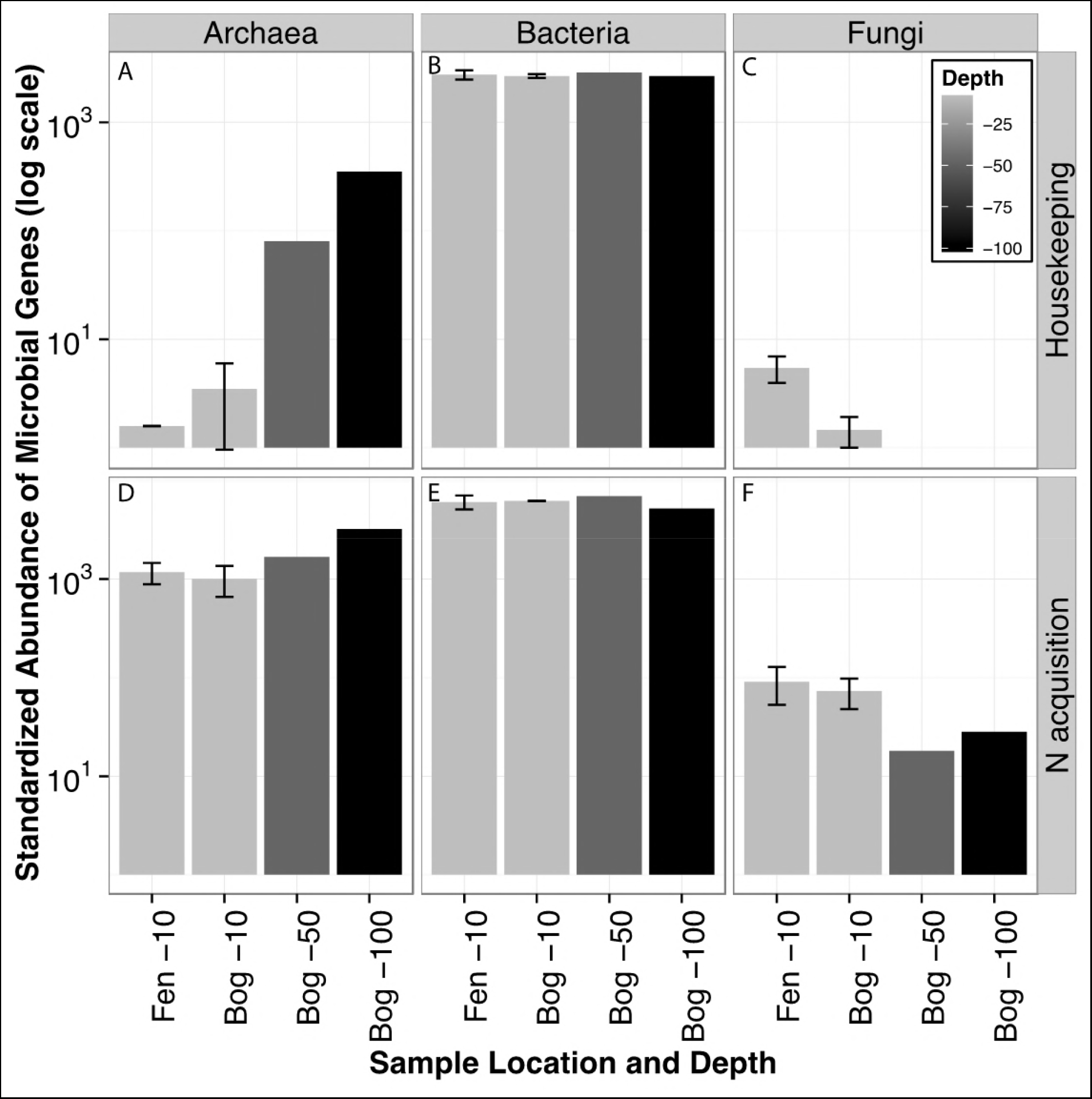
The standardized abundance of identified microbial genes in MEF peatlands through sampling depth. (A-C) show the distribution of house-keeping genes. (D-F) show the distribution of N acquisition genes. Fen −10 and Bog −10 represent the average standardized gene abundance of two surface fen and bog samples, respectively.

Few differences in standardized gene abundance were observed between bog and fen acrotelm samples, except that fungal genes were more abundant in fen acrotelm compared to bog acrotelm samples (Figure 1). At gene family level, three protease families (M14, M4_C, and Asp) were at least 12% more abundant in samples from the fen compared to bog acrotelm. Whereas assemblies resembling U56 and *nifH* genes were less abundant in the fen compared to bog. The rest of the protease genes were less than 10% different across environments (Table 2).

**Table 2.** The standardized abundance of N acquisition genes in acrotelm peat samples. The values represent the average of two replicates of acrotelm samples from each geological location. The lighter color indicates a higher value. The percent difference = 100 *(H/L-1), where H is high value and L is low value.

| Gene Categories | pfam | Average of two acrotelm replicates |  | Percent difference |
| --- | --- | --- | --- | --- |
|  |  | Bog | Fen |  |
| Protease | trypsin | 2041.332082 | 1998.470798 | 2% |
|  | S10 | 624.412566 | 643.052581 | 3% |
|  | S8 | 516.762595 | 497.759234 | 4% |
|  | M28 | 55.793862 | 59.416948 | 6% |
|  | M1 | 2614.434573 | 2823.429947 | 8% |
|  | U56 | 701.534165 | 619.595911 | 13% |
|  | M14 | 222.278808 | 251.737591 | 13% |
|  | M4_C | 7.314997 | 9.486717 | 30% |
|  | Asp | 6.773885 | 16.040373 | 137% |
| Nitrogen Fixation | *nifH* | 530.531558 | 453.149944 | 17% |

### N-acquisition genes

Genes encoding for *nifH* were detected in archaea and bacteria only, and approximately 97% of the *nifH* genes detected were bacterial. Nitrogen fixation is not known to be mediated by fungal communities (52). *NifH* genes were substantially less abundant than protease genes. The abundance of detected *nifH* genes was similar to the abundance of protease family S8, which is the least abundant archaeal protease family and the fourth least abundant bacterial protease family (Figure 2).

**Figure 2.**
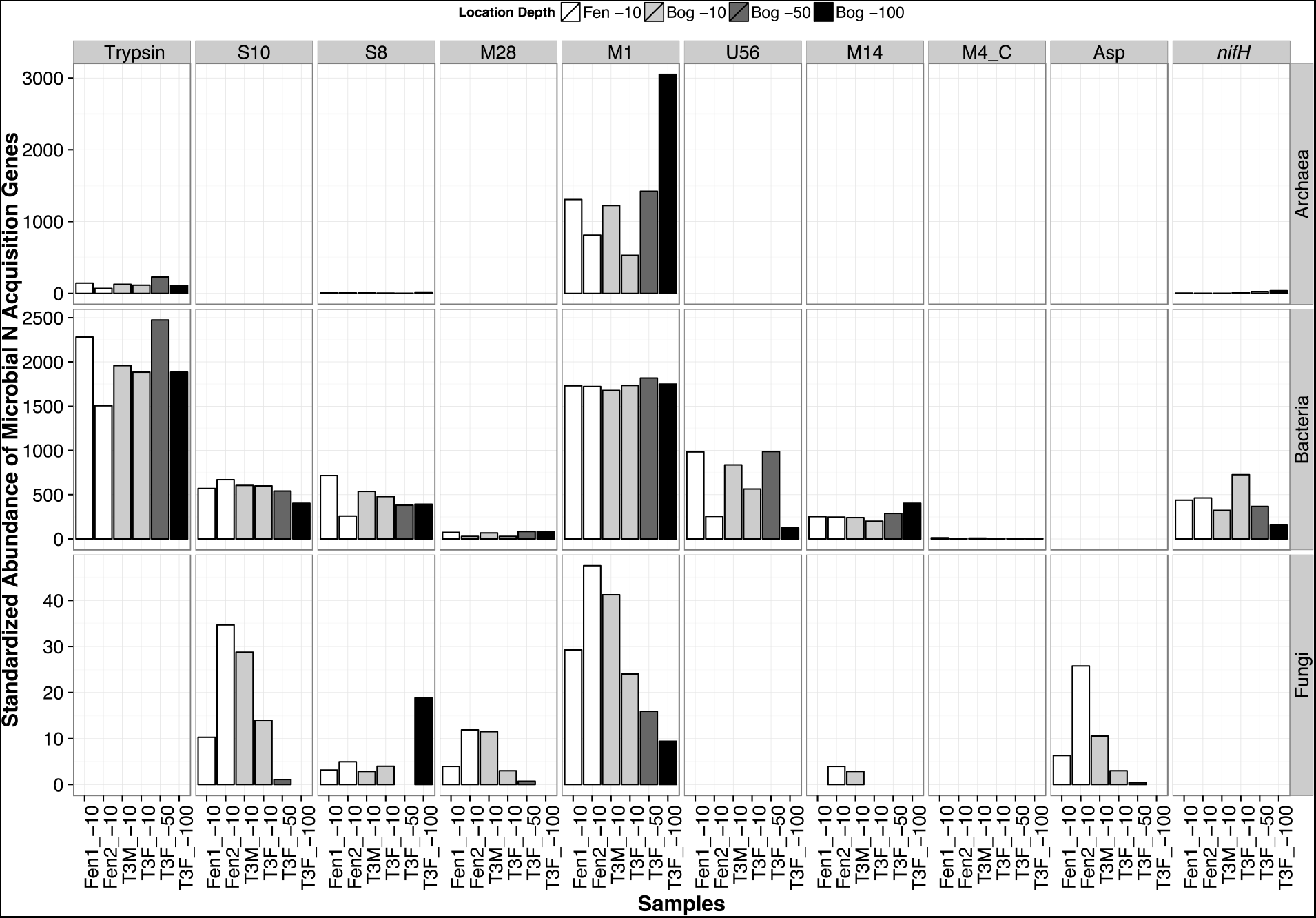
The abundance of identified nitrogen acquisition gene assemblies and their taxonomic distribution. Microbial proteases greatly outnumbered *nifH* genes (last column) with the most abundant gene families (bacterial Trypsin (column 1) and archaeal M1 (column 4) each containing more sequences than all sequences attributed to *nifH*. Additionally, the relative abundance of most bacterial protease genes did not differ across depth profiles. Samples from the same depth and environment are colored identically, as denoted in the legend. The sampling depth increases from left to right. A description of samples is located in (Table 1.

Protease encoding genes differed in distributions among archaea, bacteria, and fungi (Figure 2). Among nine protease gene families, eight were identified in bacteria, six were identified in fungi, and only three were identified in archaea. With some exceptions, archaeal protease genes increased with sampling depth, fungal protease genes decreased with depth, and bacterial protease genes varied. In contrast to these trend, archaeal Trypsin, bacterial Trypsin, M1, and U56 genes were the most abundant in the mesotelm, and fungal S8 genes were the most abundant in the catotelm and undetected in the mesotelm (Figure 2).

The protease Asp genes were uniquely detected only in fungi, while M4_C and U56 genes were uniquely detected in bacteria (Figure 2). Figure 3 shows the taxonomy distribution of the most abundant fungal Asp genes, bacterial M4_C genes, and bacterial U56 genes. In protease Asp family, a large fungal genera variation was observed among acrotelm samples. Asp genes similar to those of genera *Phanerochaete*, *Pseudogymnoascus*, and *Aspergillus* were detected in fen samples only (Figure 3A). In protease M4_C family, the bacterial genera in acrotelm sample Fen2_-10 is drastically different from the rest of the samples (Figure 3B). Protease genes similar to those found in *Methylocella* and *Burkholderia* were the most abundant genus identified with bacterial protease U56 (Figure 3C).

**Figure 3.**
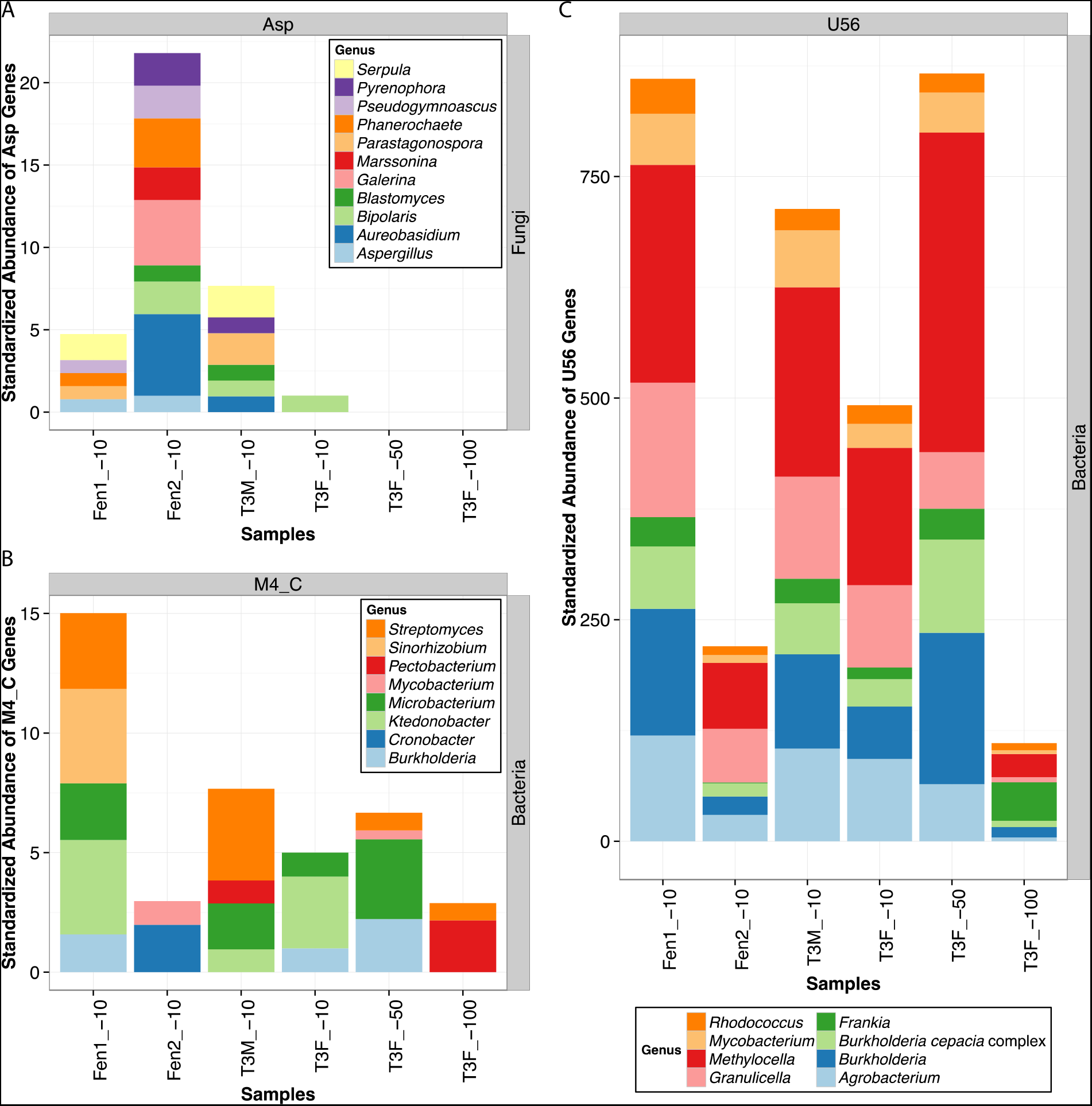
Phylogenetic distribution of the most abundant (A) fungal Asp, (B) bacterial M4_C, and (C) bacterial U56 genes with variation across environments. Genus-level data are presented.

## Discussion

Nitrogen fixation has long been considered the primary N source for peatlands (13-18), but N fixation alone cannot meet ecosystem N requirements (5, 19). Similarly, N assimilation has been shown to exceed gross mineralization in northern ecosystems (19, 20), and intact amino acid assimilation has been recognized as a potentially important source to meet N demand (22, 30, 36, 53). Previous work suggested that microbial proteases may be a missing link in northern peatland C and N cycling (30, 54). We investigated the proteolytic potential of peatland microbiomes across depth and environment type. Our work contrasts the paradigm of fungal importance in depolymerization processes and suggests that niche complementarity among diverse microorganisms is likely to play a substantial role in C and N cycling within northern peatlands.

We reveal unique niches for fungal, bacterial, and archaeal proteolytic potential, as protease families from each kingdom showed distinct stratification patterns across depth. Fungi were mostly found in the acrotelm, which constitutes the peat surface and is more oxygenated than deeper peat layers. This layer contains higher concentrations of C inputs from newly-derived plant material, such as lignin and large proteinaceous molecules. Lignin in particular requires oxygen for decomposition due to its comparatively high chemical complexity, and other work has indicated an association between fungal proteases and lignin decomposition under N-limited conditions (55, 56). Previous work in this system has also shown that carbohydrates are enriched in the surface layer, while amino sugars and saccharides increase with depth (34). We therefore suggest that fungi are particularly relevant players in early stages of decomposition, in which fresh plant material is degraded into smaller organic compounds, including proteins and oligopeptides.

Bacteria were the most abundant sequences detected regardless of depth or environment, consistent with previous work by Lin et al. (50). Both house-keeping genes and N-acquisition genes (*nifH* and protease genes) were approximately equally abundant throughout depth, and their sheer abundance suggest that at the community scale bacteria can outcompete fungi and archaea for plant-derived compounds. At gene family level, eight out of nine proteases families were detected in bacteria with a highly variable abundance across depth. Diversity throughout the depth profile despite relatively constant abundance indicates plasticity in bacterial resource use across a variety of organic matter degradation states. Additionally, large numbers of bacterial proteases relative to fungi and archaea signifies a possible dominance of bacterial depolymerization in northern peatlands.

Complementary to fungal and bacteria niche space, archaea had clear advantage in deep peat. Archaeal N-acquisition genes were consistently more abundant in the mesotelm (25-50 cm) and catotelm (>50 cm) than the acrotelm (0-10 cm) at gene family level. Lin et al. (50) noted the presence of archaea more generally at depth in peat, reaching up to 60% of total small-subunit rRNA gene sequences below 75 cm. Archaea are found in a variety of anaerobic and extreme environments and may be more tolerant of low-oxygen conditions that persist in deep peat than their bacterial or fungal counterparts. The presence of archaeal proteases at depth specifically suggests that archaea may be vital to the decomposition of the oldest and most humified organic materials stored within peatlands. In total, the consistent differences in abundance and diversity of fungi, bacteria, and archaea across the three peat depths of the bog and fen suggests that niche partitioning across redox profiles may substantially influence the mechanisms of microbial decomposition.

Protease gene abundance and taxonomic association differed between fen and bog acrotelm samples. Previous work has shown that MEF fens have roughly 10% more dissolved organic C than bogs and that this difference in geochemistry explains most of the variation in microbiome composition between fen and bog samples (47). We note that fungal protease genes in particular are more abundant in the fen acrotelm than bog acrotelm (the layer in which most fungal biomass was found), consistent with observations that fungi are more active in low-nutrient niches (57, 58). Although bacterial and archaeal protease gene abundances were mostly similar across environments, one bacterial protease gene family (U56) was 12% more abundant in the bog than the fen and mostly consisted of organisms belonging to *Methylocella* and *Burkholderia* (Table 2, Figure 3). While we did not explore the niches of these organisms beyond proteolytic activity, *Methylocella* are commonly associated with methantrophy (59) and *Burkholderia* are functionally diverse, but often considered to be plant-associated nitrogen fixers (60, 61). The abundance of proteases associated with these clades merits future investigation into their role in peatland biogeochemistry.

Regardless of environment type or depth, bacterial protease abundance and diversity as a whole indicates a wide variety of possible niches for C and N cycling bacteria within peatlands. Aminopeptidase N (M1), which cleaves peptides and produces N-terminal amino acid residues (62, 63), was the most prevalent microbial protease. Work in other systems has shown that bacterial aminopeptidase N proteases can account for 99% of alanine released from substrate hydrolysis (64) and that they are critical in generating bioavailable organic N via microbial biomass turnover (65). Thus, we highlight the M1 gene family as a key enzyme in understanding peatland N cycles.

Beyond the protease M1 family, extracellular protease genes were highly diverse. We propose that specific extracellular protease families we identified may fill unique steps in decomposition of plant material (Figure 3) (66-68). Below we discuss families that are both abundant in our samples and ecologically-relevant to peatland ecosystems. Two gene families, Asp and Aspartic endopeptidase, are commonly associated with fungal wood decomposition (55, 56). These proteases may therefore play an important role in the early stage of peatland depolymerization, in which large polymeric molecules are degraded. Asp gene families associated with *Phanerochaete*, *Pseudogymnoascus*, and *Aspergillus* were notably present only in fens (Figure 3). Proteases associated with *Phanerochaete chrysosporium* in particular are associated with highly N-limited systems (55, 56). Their high abundance within peatland metagenomes, and in the more N-limited fen environment, along with physiological selection under low N concentration reflects a distinct ecological niche for these organisms in peatlands. Finally, bacterial protease gene family U56 (formerly *linocin M18* (69)) has been largely studied within the context of dairy fermentation but is identified as a key enzyme for the decomposition of milk proteins by *Brevibacterium linens* (69). They may play a similar role within peatland by targeting proteinaceous material though further investigation is necessary.

While we also support previous work showing an the importance of N fixation in peatlands, particularly in surface peat (34), we suggest that microbial depolymerization compliments N fixation in highly N-limited ecosystems. We were able to assemble a substantial amount of archaeal and bacterial *nifH* genes (>21,000). *Sphagnum* is known to harbor a diversity of N-fixing symbionts (48), including *Cyanobacteria* observed here (70). However, *nifH* genes were much less abundant than the most abundant protease gene families (archaeal M1 and bacterial Trypsin), in line with Vile’s observation that N accumulation in boreal forests exceeded N deposition from atmosphere by 12-25 fold (10).

## Conclusion

We explored the genetic potential for organic matter depolymerization in a northern peatland based on previous work indicating that microbial proteases may be a vital uncertainty in C and N cycling within these ecosystems (30, 54). We hypothesized that i) surface peat protease genes are fungal dominated; ii) bogs contain more proteolytic potential than fens; and iii) protease gene abundance and diversity decreases with depth; however, our results were only partially consistent with these hypotheses. While we found that fungal protease genes were abundant in the acrotelm (surface layer), bacterial proteolytic potential was orders of magnitude greater and distributed through depth profiles. Prevalence of archaeal protease genes at depth suggests an importance of these organisms in C and N available below the rooting zone in peatlands. In contrast to our hypothesis, bacterial protease gene abundance was consistent across environments and fungal protease genes were more prevalent in the low-nutrient fen environment. We also show a diversity of protease genes that suggests strong niche complementarity among microorganisms with different physiologies. We identify proteases belonging to gene families M1, U56, Asp, and Aspartic endopeptidase as well as those associated with *Phanerochaete chrysosporium* as proteases that may be particularly important within northern peatlands. In total, proteases greatly outnumbered *nifH* genes attributed to N fixation, emphasizing their role in peatland C and N cycles. We contrast the historical paradigm of fungal dominance in depolymerization processes and suggest that bacteria are imperative in releasing free amino acids from peptides through depolymerization of older organic material. Our work demonstrates high genetic potential for depolymerization from a diverse suite of microorganisms beyond those typically considered, and we urge a broader perspective on the organisms mediating C and N cycles in northern peatlands.

## Materials and Methods

### Sample description

A large-scale field manipulation experiment known as Spruce and Peatland Response Under Changing Environments (SPRUCE) was initiated at the Marcell Experimental Forest (MEF), Minnesota, USA, by the U.S. Department of Energy, the U.S. Department of Agriculture (USDA) Forest Service, and Oak Ridge National Laboratory (http://mnspruce.ornl.gov/). The MEF itself is a 8.1-hectare acidic, forested bog (N47°30’31.132”, W93°27’15.146”). Sites within the MEF are classified based on their trophic status and water source as ombrotrophic bogs (receiving precipitation only) or minerotrophic fens [fed by both groundwater and precipitation (47, 71)]. Although fens are frequently considered more nutrient rich than bogs, both types of peatlands are highly limited in inorganic N. A full characterization of the field site including peatland hydrology and vegetation is described by Sebestyen et al. (72). Further information on samples is available in Lin et al. (34, 50).

Six metagenomic libraries were obtained from MEF in February 2012 as per Lin *et al*. (34). Briefly, peat cores were collected from hollows in bogs and fens and sectioned from 0- to 10- (acrotelm), 25- to 50- (mesotelm), and 75- to 100-cm (catotelm). The water table was at the surface of the *Sphagnum* layer. Each core section was homogenized. Two acrotelm samples (0 to −10 cm) were collected from bog lake fen (i.e. samples Fen1_-10 and Fen2_-10), two acrotelm samples (0 to −10 cm) from SPRUCE bog (i.e. samples T3M_-10 and T3F_-10), one mesotelm sample (−25 to −50 cm) from SPRUCE bog (i.e. sample T3F_-50), and one catotelm sample (−75 to −100 cm) from SPRUCE bog (i.e. sample T3F_−100). As described in Lin et al. (34, 50), samples differed in physicochemical properties across the depth layers. Sequencing coverage for each metagenome increased with depth and ranged from 42% to 86% (Table 1). Metagenomes were generally consistent between samples from the same depth and site, suggesting that the results reported here are likely to be robust (further details provided in Lin et al. (34)). Whole genome shotgun metagenome sequences are available in MG-RAST (34, 50).

### Hidden Markov Model construction

We used 24 Hidden Markov Models (HMMs) constructed based on protein sequences (73) to investigate the microbial genetic potential in N acquisition in MEF peatlands, 20 of which were for microbial protease genes, one for nitrogenase gene (*nifH*), and three for microbial single copy housekeeping genes (Table S2). Nitrogenase enzymes are encoded by three genes *[nifH, nifD*, and *nifK* (reviewed in 74)]; however *nifH* is the most commonly used marker gene for nitrogenase potential (52, 75). Evidence suggests that *nifD-* and nifK-based assays are consistent with *nifH-based* results (76). Hereafter, we use ‘N-acquisition genes’ and related terms to represent all protease genes and *nifH*. The abundance of N-acquisition genes is therefore the sum of the abundance of all protease genes plus the abundance of *nifH*. The housekeeping genes used as bacterial, fungal, and archaeal markers are genes encoding for ribosomal protein L2 (*rplB*), RNA polymerase second largest subunit domain 4 (RPB2_4), and domain 7 (RPB2_7), respectively. We use the term ‘housekeeping genes’ to represent these domain-specific single copy housekeeping genes. The abundance of housekeeping genes is calculated as the sum of the abundance all housekeeping genes. The HMM’s for *nifH* and *rplB* genes are available in Ribosomal Database Project (RDP) Fungene repository (77). Models for RPB2_4 and RPB2_7 were obtained from Pfam database (http://pfam.sanger.ac.uk/). The protease genes (one intracellular and 19 extracellular) were selected based on literature characterizations (78-80). To construct the most informative HMM models for targeted gene groups, well-studied gene sequences were first selected from existing literature. These seed sequences were cross-checked against the reviewed protein database SwissProt (http://www.uniprot.org/). Genes encoding for proteases were also searched against the MEROPS peptidase database (http://merops.sanger.ac.uk/) to confirm their protease identity. Protein families and existing protein HMM models were queried from Pfam database. The retrieved Pfam HMMs were then used to extrapolate archaeal, bacterial, and fungal reference protein sequences from UniProt database. Pfam HMMs were used to search against SwissProt at different cutoffs (E value or Bit score) to ensure model accuracy. If existing Pfam HMMs could not accurately query sequences, a set of well-annotated sequences would be used to construct new models (Table S2). Reference protein sequences were retrieved from UniPort and aligned using finalized HMMs.

### Guided metagenomic assembly

All metagenomic reads were filtered by using RDP SeqFilters (81) to a minimal average read quality of Q = 25. Genes with HMM models (Table S2) were assembled from combined filtered reads by using modified RDP Xander skeleton analysis pipeline (https://github.com/fishjord/xander_analysis_skel). Briefly, a De Brujin graph is built for the combined shotgun metagenome dataset. Potential gene start points (Kmer starts, k = 30 nucleotides) were identified from each gene reference sequences. Local assembling was carried out by searching constructed De Bruijin graphs at the given gene start points. These local assemblages were then merged to form the longest contigs possible. The final merged nucleotide sequences were dereplicated using CD-Hit 4.6.1 (-c 1.0) (82) to identify the longest unique contigs.

### Data processing

All quality filtered reads were mapped (Bowtie2.2.5)(83) against the dereplicated merged contigs. Only contigs that were 100% covered (median base coverage = 1) were considered. The biological information of these fully covered contigs were identified using Basic Alignment Searching Tool (blastx) 2.2.30+; the best matching sequences with E-values ≤ 1×10^−5^ were kept (84). UniProt (UniProtKB release 10, 2014) was used as the annotation database.

Final gene abundance per peat sample was determined by mapping reads from each metagenome to fully covered contigs. The mapping results indicated ~9% of reads were mapped onto the contigs once only and majority of the fungal housekeeping contigs were mapped once. Hence, for downstream analyses, we included all reads mapped to final fully covered contigs at least once (Table S3). The mapped read abundances were standardized by sequencing depth for comparisons among samples. Gene abundance will be used to infer the abundance of mapped reads to fully covered contigs from now on unless otherwise specified. Final data analyses and visualization was done in R 3.1.0 (85) with packages plyr (86) and ggplot2 (87).

## Acknowledgements

We would like to thank Dr. Joel E. Kostka (Georgia Institute of Technology) for providing the metagenome sequences (collaboration with Dr. Christopher W. Schadt, Oak Ridge National Laboratory) and the Ribosomal Database Project (Michigan State University) for their help on using Xander analysis skeleton pipeline. This research is supported by the DOE Office of Science (Office of Basic Energy Sciences), grant no. ER65430.

